# Appetitive Pavlovian goal-tracking memory reconsolidation is reduced by both adrenergic and NMDA receptor antagonism

**DOI:** 10.64898/2026.06.23.733991

**Authors:** Jonathan L. C. Lee

## Abstract

**Rationale:** Appetitive Pavlovian cues can drive maladaptive reward seeking via stimulus–reward memories. Disrupting memory reconsolidation offers a potential strategy to reduce their influence, but evidence for β-adrenergic blockade with propranolol is inconsistent across behavioural paradigms, particularly relative to NMDA receptor antagonism.

**Objectives:** We tested whether propranolol disrupts reconsolidation of appetitive sucrose memories in a discriminative goal-tracking paradigm, and compared its effects with those of the most commonly used NMDA receptor antagonist, MK-801.

**Methods:** Adult Lister hooded rats underwent discriminative Pavlovian conditioning. Thirty minutes before a brief memory reminder (non-reinforced or reinforced), rats received systemic drug treatment or saline control. In study 1, MK-801 (0.1 mg/kg) was administered to male rats. In study 2, propranolol (10 mg/kg) was administered to equal numbers of male and female rats. Goal-tracking was tested drug-free at 1 and 8 days.

**Results:** In study 1, MK-801 impaired subsequent discriminated responding at test. These effects were observed not only when reminder was non-reinforced as in previous successful demonstrations, but also with reinforced reminder. In study 2, Propranolol also impaired subsequent goal-tracking, regardless of reminder type, and the effects were consistent across sexes.

**Conclusions:** Propranolol can disrupt reconsolidation of appetitive goal-tracking memories to a similar extent as MK-801 under conditions that promote memory destabilisation. These findings demonstrate that β-adrenergic blockade can impair appetitive memory reconsolidation in a goal-tracking paradigm, challenging prior null findings and revitalising the potential for propranolol-based interventions in maladaptive reward-seeking behaviours.

## Introduction

Stimuli that are predictive of rewarding outcomes can gain appetitive incentive properties that can manifest in a range of behavioural settings. For example, appetitive pavlovian stimuli can act as reinforcers in their own right, invigorate reward-seeking behaviour, and draw attention and behaviour towards themselves (Everitt et al. 2003). Each of these behavioural impacts of appetitive pavlovian stimuli may contribute in important ways to pathological reward-seeking behaviours (e.g. in drug addiction)(Milton and Everitt 2012).

Within this context, efforts to diminish the strength and impact of stimulus-reward associations have been undertaken, with one area of focus being the impairment of memory reconsolidation (Milton and Everitt 2010; Barak and Goltseker 2021; Milton 2023). Reconsolidation impairments have been observed in conditioned reinforcement, pavlovian-instrumental transfer and autoshaping/sign-tracking paradigms, using both natural rewards and addictive drug rewards (Lee and Everitt 2008a, b; Lee et al. 2005; Milton et al. 2008b, a, 2011). Among the variety of amnestic agents used in such studies, commonly-used systemically-administered drugs are the NMDA receptor antagonist MK-801and the adrenergic receptor antagonist propranolol. While both drugs impair conditioned reinforcement within an acquisition of a new response paradigm with sucrose and cocaine reward (Milton et al. 2008b), their effects diverge in pavlovian-instrumental transfer and autoshaping settings, with only MK-801 (and not propranolol) producing impairments with sucrose and ethanol rewards (Lee and Everitt 2008a; Milton et al. 2011). This raises the possibility that MK-801 and propranolol will not have equivalent disruptive effects in attempts to impair appetitive memory reconsolidation in other settings.

Other commonly-used appetitive memory paradigms are conditioned place preference and pavlovian conditioned approach (also called goal-tracking). While several studies have shown reconsolidation impairments following propranolol and MK-801 treatments in conditioned place preference for a variety of natural and addictive drug rewards (Bernardi et al. 2006; Kelley et al. 2007; Brown et al. 2008; Fricks-Gleason and Marshall 2008; Robinson et al. 2010; Achterberg et al. 2012, 2014; Alaghband and Marshall 2012; Liu et al. 2015), there has been much less exploration in goal-tracking paradigms.

Goal-tracking captures cue-driven, outcome-directed behaviour mediated by explicit or implicit outcome expectancy, and is a core component of human cue-reactivity in addiction and appetitive disorders (Savage and Ramos 2009; Morrison et al. 2015; Lovibond and Westbrook 2025). Early studies in goal-tracking settings showed little evidence of impairments in memory reconsolidation, which suggested a resistance of outcome-directed behaviour to reconsolidation-based interventions. While the systemic administration of anisomycin immediately after conditioning sessions impaired the acquisition of a goal-tracking pavlovian conditioned approach response, the same treatment after memory reminder had no obvious disruptive effect on previously-established goal tracking (Blaiss and Janak 2007). A similar pattern of post-training modulation, without post-reminder effects, was observed with systemic amphetamine injections (Blaiss and Janak 2007). In contrast, we have previously demonstrated that under specific parametric conditions of conditioning and reminder, systemic injections of MK-801 prior to memory reminder do impair memory reconsolidation in a goal-tracking setting (Reichelt and Lee 2012). Similarly, post-reminder extinction or fear conditioning impaired goal-tracking in a subset of rats (those that orient to the conditioned stimulus; (Olshavsky et al. 2013)).

In this context of somewhat inconsistent findings, a recent study using systemic injections showed that post-reminder propranolol did not impair goal-tracking responses, whereas it did impair sign-tracking responses in different animals (Cogan et al. 2019). While the contrast between the impaired sign-tracking and preserved goal-tracking is certainly interesting, especially as the two observations resulted from parametrically-aligned experiments, considering the goal-tracking experiment in isolation suggests at least two potential accounts of the negative effect. First, the reminder consisted of sessions that were identical to the training sessions; at the theoretical level, it is suggested that prediction error or mismatch at reminder is important to trigger memory reconsolidation (Pedreira et al. 2004; Exton-McGuinness et al. 2015). Second, and somewhat related, the conditioning phase may have resulted in near-asymptotic levels of learning; we and others have shown that reconsolidation impairments are more reliably observed under conditions of prior sub-asymptotic learning (Rodriguez-Ortiz et al. 2005, 2008; Reichelt and Lee 2012, 2013), including when reminder consists of an additional learning experience (Rodriguez-Ortiz et al. 2005, 2008; Lee 2008). Here, we tested whether systemic administration of propranolol impairs memory reconsolidation to reduce discriminative goal-tracking behaviour, benchmarking its effects against the established efficacy of MK-801 (Reichelt and Lee 2012), with the aim of clarifying the translational potential of adrenergic receptor-based reconsolidation interventions for maladaptive cue-driven reward seeking.

## Methods

### Subjects

The subjects were 71 experimentally-naïve adult Lister hooded rats (Charles River, UK). Male rats weighed 200-250 g at the start of the experiment; female rats weighed 175-200 g at the start of the experiment. Rats were housed in groups of four in standard cages. Holding room conditions were 21°C and a standard 12-hr light/dark cycle (lights on at 0700). Food and water were available ad libitum in the holding cages. Training and testing were conducted between 1000 and 1500. All procedures were conducted in accordance with the United Kingdom 1986 Animals (Scientific Procedures) Act (Amendment Regulations 2012)(PPLs 70/7662 & PP7563911).

### Drugs

The rats were administered (+)-5-methyl-10,11-dihydro-5Hdibenzo[a, d]cyclohepten-5,10-imine maleate (MK-801, Sigma) dissolved in saline (0.1 mg/mL) at a dose of 0.1 mg/kg, or DL-propranolol hydrochloride (Sigma) also dissolved in saline (10 mg/mL) at a dose of 10 mg/kg. Injections were administered intraperitoneally (i.p.) 30 min prior to the reminder session, and doses were identical to those used previously to impair appetitive memory reconsolidation (Lee and Everitt 2008a; Reichelt and Lee 2012). Saline served as the vehicle control.

### Design

For each study, drug administration was compared against vehicle control, with individual rats constituting the experimental unit. The target sample size for each experimental group was 6-8, based upon previous studies and an expected large effect size.

Study 1: MK-801 or saline was administered. Only male rats were used. MK-801 non-reinforced reminder groups had 7 animals and reinforced reminder groups had 6 animals in each condition.

Study 2: Propranolol or saline was administered. The propranolol experiments were pre-registered (https://osf.io/92hgm/) and equal numbers of male and female rats were used. All propranolol groups had 8 animals.

Experiments were run separately and rats were allocated to group within each experiment at random. Where sub-cohorts were run, randomisation was applied at the sub-cohort level, taking into account any requirement to balance numbers. Randomisation was achieved using the sequence function at Random.org. No additional approaches were taken to avoid potential confounds. The truly-random approach did not result in any occurrences of all animals in a single cage being allocated to the same group.

Experimenters were not strictly blind to experimental group at any stage of the study. However, when handling and injecting rats, group allocation was concealed.

### Behavioural procedures

All procedures took place in the same experimental room, within operant chambers as described previously (MedAssociates; (Reichelt and Lee 2012)). The behavioural procedures were also identical to those used previously (Reichelt and Lee 2012).

### Conditioning

Pavlovian conditioning was carried out 6 days (one session per day; Tuesday-Friday, Monday-Tuesday) during which two discriminable auditory stimuli (click or tone) were presented 10 times each session for 30 sec. One stimulus (the CS+, counterbalanced across the rats) was reinforced with the delivery of three sucrose pellets (45 mg; P.J. Noyes) at the end of the presentation, whereas the CS – was never reinforced. Each stimulus presentation was separated by a 60-sec period, separated into a 30-sec intertrial interval (ITI) and a 30-sec PreCS period, in which no cues were presented.

### Memory reminder

Memory reminder took place the day after the end of conditioning. Drug or vehicle was administered 30 min prior to the reminder procedure. There were two reminder conditions. The non-reinforced reminder consisted of a short extinction session where three presentations each of the CS+ and CS– were presented without being rewarded. Each stimulus presentation was separated by a 60-sec period, separated into a 30-sec ITI and a 30-sec PreCS period, in which no cues were presented. This was operationally identical to conditioning sessions, but with no sucrose reward and only 3 presentations of each CS. The reinforced reminder was the same, but with each CS+ presentation being reinforced by 3 sucrose pellets as in the conditioning sessions.

### Test

Test sessions were conducted one day after memory reminder and additionally 7 days later. At test, goal-tracking activity was measured by magazine entry behaviours during non-reinforced CS+ and CS- presentations, and thus the clicks or tones were presented 10 times for 30 sec, each trial separated by a 60-sec stimulus-free period, separated into a 30-sec ITI and a 30-sec PreCS period.

### Measures and inclusion criteria

Approach ratios for the CS+ and CS- were calculated for conditioning, memory reminder and test sessions. This was measured by ratios of magazine responding during the CS presentations and their preceding periods as calculated by the equation CS/(CS + PreCS). The primary outcome measure was the discrimination in approach ratios at the test sessions (i.e. the comparison of approach to the CS+ and CS- between groups).

Only rats that showed goal tracking behaviour at the end of conditioning were included in the analyses. This was determined by *a priori* criteria of the CS+ approach ratio being >0.5, and the CS+ approach ratio being greater than the CS- approach ratio. i.e. rats approached the magazine more during CS+ presentation than during CS- presentation, and also more during CS+ presentation than the preceding baseline PreCS period.

Approach ratios were calculated only after completion of all procedures. 3 rats were excluded from the MK-801 non-reinforced reminder experiment (2 x Saline; 1 x MK-801). 4 rats were excluded from the MK-801 non-reinforced reminder experiment (2 x Saline; 2 x MK-801). No rats were excluded from either of the propranolol experiments. There was no missing data relevant to the statistical analyses (there was some missing data from the first day of conditioning).

### Statistical analyses

Data were analysed in JAMOVI (The jamovi project 2025). The approach was used in the MK-801 experiments and then pre-registered prior to the propranolol experiments. The analyses described are those conducted for the MK-801 experiments. The additional factor of Sex was included for all propranolol analyses. Alpha was set at 0.05, and η_p_^2^ was reported as a measure of effect size.

For the test sessions, 2 x 2 x 2 mixed ANOVAs with factors CS, Group and Session were conducted. 2 x 2 mixed ANOVAs were conducted for the final day of conditioning and the reminder session (separately). Where significant interactions were observed, analyses of simple main effects were carried out.

We also conducted overall analyses, including reminder method as an additional factor. This was in anticipation that the method of reminder would not be significantly important. As the MK-801 and propranolol experiments were carried out at different times, no direct statistical comparison between the two was conducted.

## Results

### Study 1: MK-801

Rats acquired discriminated approach to the CS+ (data not shown). On the final day of conditioning, there was a significantly greater approach ratio to the CS+ than to the CS- (non-reinforced reminder: F(1,12)=28.3, p<0.001, η_p_^2^=0.702; reinforced reminder: F(1,10)=20.2, p=0.001, η_p_^2^=0.669). In the non-reinforced reminder condition, there was a CS x Group interaction (F(1,12)=5.12, p=0.043, η_p_^2^=0.299), driven by higher approach to the CS- in the saline group than the MK-801 group. In contrast, there were no differences between the groups in the reinforced reminder condition (CS x Group: F(1,10)=0.704, p=0.421, η_p_^2^=0.066; Group: F(1,10)=1.80, p=0.210, η_p_^2^=0.152).

At memory reminder, there was no significant acute effect of MK-801 on approach behaviour in the non-reinforced reminder condition (data not shown; CS x Group: F(1,12)=0.0429, p=0.839, η_p_^2^=0.004; Group: F(1,12)=0.274, p=0.610, η_p_^2^=0.022). There was, however, an acute effect of MK-801 to impair discriminated approach in the reinforced reminder condition (CS x Group: F(1,10)=5.31, p=0.044, η_p_^2^=0.347; no effect of group on CS+ or CS- approach ratios individually).

At test, there was evidence for an effect of MK-801 to impair discriminated responding across both tests. In the non-reinforced reminder condition (Fig. 1A), there was a CS x Group interaction (F(1,12)=7.20, p=0.020, η_p_^2^=0.375), with no CS x Group x Session interaction (F(1,12)=2.26, p=0.159, η_p_^2^=0.158). Analyses of simple main effects of Group revealed an MK-801 impairment on CS+ approach ratios (Group: F(1,12)=9.79, p=0.009, η_p_^2^=0.449; Test x Group: F(1,12)=0.293, p=0.598, η_p_^2^=0.030), but not on CS- approach ratios (Group: F(1,12)=0.027, p=0.871, η_p_^2^=0.002; Test x Group: F(1,12)=2.24, p=0.160, η_p_^2^=0.158). Orthogonal simple main effects revealed discriminated approach in Saline rats (CS: F(1,6)=17.9, p=0.006, η_p_^2^=0.748; Test x CS: F(1,6)=1.71, p=0.239, η_p_^2^=0.221), but not in MK-801 rats (CS: F(1,6)=0.289, p=0.610, η_p_^2^=0.046; Test x CS: F(1,6)=0.711, p=0.431, η_p_^2^=0.106).

**Figure 1.**
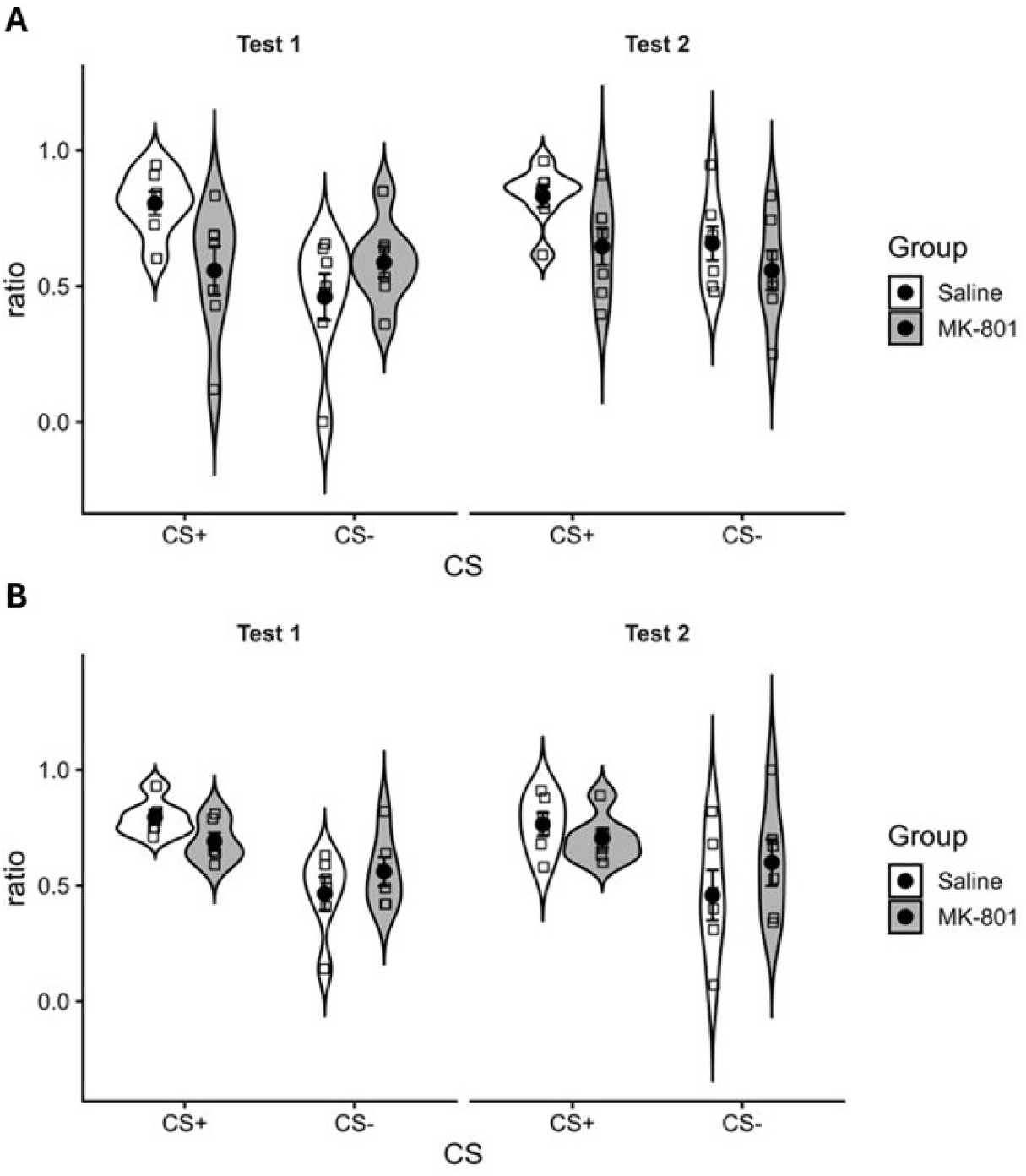
Pre-reminder MK-801 impairs subsequent discriminative goal-tracking. MK-801 was injected prior to non-reinforced (**B**) or reinforced (**A**) reminder. Discriminative approach to the reward location was tested 1 (Test 1) and 8 days (Test 2) later. MK-801-treated rats showed reduced discriminative goal-tracking across tests and for both reminder types. Summary data presented as mean ± SEM.

In the reinforced reminder condition (Fig. 1B), there was a CS x Group interaction (F(1,10)=14.6, p=0.003, η_p_^2^=0.594), with no CS x Group x Session interaction (F(1,10)=5.43 x 10^-4^, p=0.982, η_p_^2^=0.000). Analyses of simple main effects of Group revealed no clear MK-801 impairment on CS+ approach ratios (Group: F(1,10)=3.71, p=0.083, η_p_^2^=0.270; Test x Group F(1,10)=0.304, p=0.593, η_p_^2^=0.030), or CS- approach ratios (Group: F(1,10)=1.97, p=0.191, η_p_^2^=0.164; Test x Group: F(1,10)=0.0642, p=0.805, η_p_^2^=0.006). Orthogonal simple main effects revealed discriminated approach both in Saline rats (CS: F(1,5)=135.2, p<0.001, η_p_^2^=0.964; Test x CS: F(1,5)=0.048, p=0.836, η_p_^2^=0.009), and in MK-801 rats (CS: F(1,5)=7.05, p=0.045, η_p_^2^=0.585; Test x CS: F(1,5)=0.079, p=0.791, η_p_^2^=0.015).

Analysing both reminder conditions together revealed no obvious dependence of MK-801 effect upon the nature of the reminder. There remained a CS x Group interaction (F(1,20)=18.6, p<0.001, η_p_ ^2^=0.481), with no CS x Group x Session interaction (F(1,20)=1.04, p=0.319, η_p_ ^2^=0.050), no CS x Group x Reminder interaction (F(1,20)=0.123, p=0.730, η_p_ ^2^=0.006), and no CS x Group x Session x Reminder interaction (F(1,20)=1.08, p=0.311, η_p_ ^2^=0.051).

### Study 2: Propranolol

Rats acquired discriminated approach to the CS+ (data not shown). On the final day of conditioning, there was a significantly greater approach ratio to the CS+ than to the CS- (non-reinforced reminder: F(1,12)=68.0, p<0.001, η_p_^2^=0.850; reinforced reminder: F(1,12)=47.5, p<0.001, η_p_^2=^0.798). In the non-reinforced reminder condition, there were no differences between the groups (CS x Group: F(1,12)=0.807, p=0.387, η_p_^2=^0.063; CS x Group x Sex: F(1,12)=0.010, p=0.922, η_p_^2=^0.001). In contrast, there was a CS x Group interaction in the reinforced reminder condition (F(1,12)=8.10, p=0.015, η_p_^2^=0.403; CS x Group x Sex: F(1,12)=1.10, p=0.315, η_p_^2^=0.084), which was not driven by any differences between the groups in approach ratios to the CS+ or the CS-, and there was discriminated responding in both groups. Visually, the interaction did reflect lesser discrimination in the propranolol group than in the Saline group.

At memory reminder, there was no significant acute effect of propranolol on approach behaviour in either reminder condition (data not shown; non-reinforced reminder - CS x Group x Sex: F(1,12)=0.008, p=0.930, η_p_^2^=0.001; CS x Group: F(1,12)=0.274, p=0.610, η_p_^2^=0.022; Group: F(1,12)=0.574, p=0.463, η_p_^2^=0.046; reinforced reminder - CS x Group x Sex: F(1,12)=0.236, p=0.636, η_p_^2^=0.019; CS x Group: F(1,12)=0.377, p=0.551, η_p_^2^=0.030; Group: F(1,12)=0.063, p=0.807, η_p_^2^=0.005). The lack of group effect in the reinforced reminder condition suggests that the previous difference at the end of conditioning did not reflect genuine differences in conditioning.

At test, there was evidence for an effect of propranolol to impair discriminated responding across both tests. In the non-reinforced reminder condition (Fig. 2A), there was a CS x Group interaction (F(1,12)=6.39, p=0.027, η_p_^2^=0.347), with no CS x Group x Session (F(1,12)=0.423, p=0.528, η_p_^2^=0.034), CS x Group x Sex (F(1,12)=0.270, p=0.613, η_p_^2^=0.022) or CS x Group x Session x Sex (F(1,12)=0.946, p=0.350, η_p_^2^=0.073) interactions. Analyses of simple main effects of Group did not support a propranolol impairment on CS+ approach ratios (Group: F(1,12)=1.41, p=0.258, η_p_^2^=0.105; Test x Group: F(1,12)=0.117, p=0.739, η_p_^2^=0.010), but did reveal an effect to increase CS- approach ratios (Group: F(1,12)=11.5, p=0.005, η_p_^2^=0.489; Test x Group: F(1,12)=0.874, p=0.368, η_p_^2^=0.068). Orthogonal simple main effects revealed discriminated approach in Saline rats (CS: F(1,6)=26.2, p=0.002, η_p_^2^=0.814; Test x CS: F(1,6)=0.011, p=0.919, η_p_^2^=0.002), but not in propranolol rats (CS: F(1,6)=0.245, p=0.638, η_p_^2^=0.039; Test x CS: F(1,6)=0.669, p=0.445, η_p_^2^=0.100).

**Figure 2.**
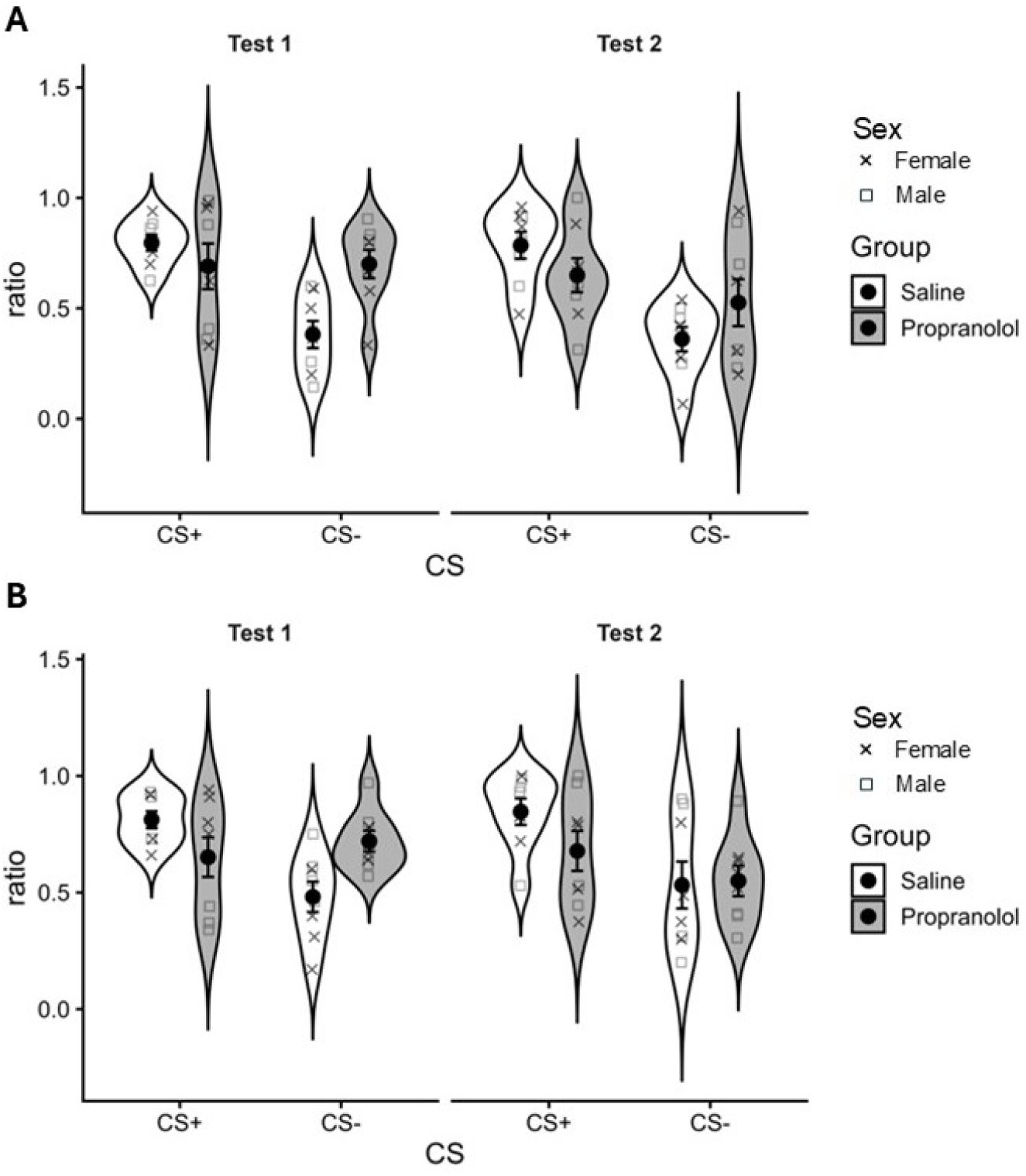
Pre-reminder propranolol impairs subsequent discriminative goal-tracking. Propranolol was injected prior to non-reinforced (**B**) or reinforced (**A**) reminder. Discriminative approach to the reward location was tested 1 (Test 1) and 8 days (Test 2) later. Propranolol-treated rats showed reduced discriminative goal-tracking across tests, regardless of rat sex, and for both reminder types. Summary data presented as mean ± SEM.

In the reinforced reminder condition (Fig. 2B), there was a CS x Group interaction (F(1,12)=6.11, p=0.011, η_p_^2^=0.427), with no CS x Group x Session (F(1,12)=0.171, p=0.215, η_p_^2^=0.125), CS x Group x Sex (F(1,12)=0.001, p=0.975, η_p_^2^=0.000) or CS x Group x Session x Sex (F(1,12)=2.86, p=0.117, η_p_^2^=0.192) interactions. Analyses of simple main effects of Group revealed a propranolol impairment on CS+ approach ratios (Group: F(1,12)=5.57, p=0.036, η_p_^2^=0.317; Test x Group F(1,12)=0.0055, p=0.942, η_p_^2^=0.000), with no effect on CS- approach ratios (Group: F(1,12)=2.33, p=0.153, η_p_^2^=0.162; Test x Group: F(1,12)=3.33, p=0.093, η_p_^2^=0.217). Orthogonal simple main effects revealed discriminated approach in Saline rats (CS: F(1,6)=14.4, p=0.009, η_p_ ^2^=0.706; Test x CS: F(1,6)=0.016, p=0.903, η_p_ ^2^=0.003), but not in propranolol rats (CS: F(1,6)=0.139, p=0.722, η_p_ ^2^=0.023; Test x CS: F(1,6)=3.72, p=0.102, η_p_ ^2^=0.383).

Analysing both reminder conditions together revealed no obvious dependence of propranolol effect upon the nature of the reminder. There remained a CS x Group interaction (F(1,24)=12.4, p=0.002, η_p_ ^2^=0.341), with no CS x Group x Session interaction (F(1,24)=1.83, p=0.188, η_p_ ^2^=0.071), no CS x Group x Reminder interaction (F(1,24)=0.140, p=0.712, η_p_ ^2^=0.006), and no CS x Group x Session x Reminder interaction (F(1,24)=0.142, p=0.709, η_p_ ^2^=0.006). No interactions with the Sex factor were significant.

## Discussion

In the present study, we observed that pre-reminder injections of MK-801 resulted in impaired discriminative goal-tracking performance across 2 tests (1 and 8 days after reminder). This occurred regardless of whether the reminder constituted non-reinforced or reinforced re-exposure to the CS+ (alongside continued non-reinforced re-exposure to the CS-). A similar pattern of results was observed with pre-reminder injections of propranolol. Therefore MK-801 and propranolol treatment have seemingly equivalent effects on the reconsolidation of appetitive memory when tested in a goal-tracking setting.

Traditionally, demonstrations of memory reconsolidation impairments have met various criteria (Dudai 2004), not all of which have been implemented in the current study. Here we show long-lasting impairments at tests both 1 and 8 days after reminder and drug treatment, which is consistent with longer-term memory impairment, rather than temporary retrieval failure. We do not have a no-reminder (or non-reactivation) control condition, and so cannot strictly rule out the possibility that drug treatment causes long-lasting effects on goal-tracking performance in a manner that does not require the putative reminder-induced destabilisation of the pre-existing memory. However, previous studies of memory reconsolidation impairments in appetitive settings using both MK-801 and propranolol have consistently demonstrated clear reactivation-dependence (Milton et al. 2008b; Lee and Everitt 2008a). Therefore, it is reasonable to conclude that the present drug-induced impairments are highly likely to be dependent upon the memory reminder, especially so for MK-801 where we have previously demonstrated reactivation-dependence (Reichelt and Lee 2012).

A final recommendation for reconsolidation demonstrations is for drug treatment to be applied after memory reminder and/or for there to be a test of post-reactivation short-term memory (Dudai 2004). The latter aligns with the theoretical assumption that memory reconsolidation relates to the long-term restabilisation of the destabilised memory, and is not necessary for short-term memory expression following memory reactivation (Nader 2003), which is presumably supported by similar mechanisms to post-learning short-term memory. Appetitive memory reconsolidation studies do not typically include post-reactivation short-term memory tests (apart from some conditioned place preference studies), at least in part due to the relatively rapid extinction of conditioned responding. Moreover, while it would be useful in future to test whether post-reactivation administration of MK-801 or propranolol is equally effective in impairing goal-tracking performance, pre-reactivation treatment ensures that pharmacological effects are more likely to be active during any peri-reactivation critical window of memory vulnerability. Nevertheless, pre-reactivation treatment does allow for acute effects during the reminder session that themselves might account for longer-term changes in performance in a manner that need not invoke memory reconsolidation processes (e.g. direct association between drug effects and the memory or appetitive cue).

Our present demonstration that pre-reminder MK-801injection impairs subsequent goal-tracking at test replicates and extends our previous findings (Reichelt and Lee 2012; Drame et al. 2020). Previously, we showed that reconsolidation effects were observed using a non-reinforced reminder procedure, but only under specific parametric conditions of training and reminder that were used in the present study (Reichelt and Lee 2012). Here we show that the goal-tracking impairments persist for at least 8 days, compared to the singular 1-day test previously. We also show that a reinforced reminder is equally effective at destabilising the underlying appetitive memory. Reinforced reminders, while certainly less commonly used in reconsolidation studies than non-reinforced reminders, have previously been shown to be effective at destabilising memory, particularly when previous learning is non-asymptotic (Rodriguez-Ortiz et al. 2005, 2008; Lee 2008). Based on our previous experiments, the 6 days of training used here similarly does not result in asymptotic learning (Reichelt and Lee 2012). While the impairments at test were clear and consistent across reminder procedures, there were somewhat unexpected statistical differences between the MK-801 and Saline groups during training for the non-reinforced reminder and at the reinforced reminder. For the training difference, this was prior to any drug treatment and showed quantitatively smaller goal-tracking behaviour in the Saline group. Therefore, it is unlikely to have had any bearing on the later impairment following MK-801 treatment. For the acute effect of MK-801 on goal-tracking behaviour during the reinforced reminder, it is possible that this contributes to the later impairment at test (either directly or by indicating that goal-tracking performance is somehow less persistent in these rats irrespective of the drug treatment). However, the quantitative impairment at test is more marked at the test than at reminder, which remains consistent with a memory reconsolidation impairment.

The present demonstration that pre-reminder propranolol impairs subsequent goal-tracking behaviour is novel and contrasts importantly with two previous studies. First, a prior study with similar food pellet reward did not show any impairing effect of propranolol on goal-tracking behaviour (Cogan et al. 2019). There are several important differences between our present study and that of Cogan et al (2019). First, Cogan et al (2019) used post-reminder injections of propranolol. While it is possible that this resulted in the pharmacological effects of propranolol falling outside the critical window of memory vulnerability, such an interpretation is weakened by the fact that the same treatment did impair sign-tracking behaviour within the same study.

A second difference is that our study used training parameters designed to achieve sub-asymptotic learning (Reichelt and Lee 2012), whereas visual inspection of the Cogan et al (2019) data indicates that learning was asymptotic prior to reminder and treatment, regardless of whether goal tracking was conditioned to a visual or auditory cue. As such, the use of a full training session as a reminder may not have created the conditions necessary to destabilise the memory via mismatch or prediction error processes. One possibility is that this factor is particularly important for goal-tracking behaviour. In contrast to sign-tracking, which is relatively insensitive to outcome value and is often considered to reflect the attribution of incentive salience to the cue, goal-tracking depends upon a representation of the predicted outcome and is sensitive to outcome devaluation (Morrison et al. 2015). Therefore, goal-tracking may rely on a more flexible, expectancy-based stimulus–outcome representation that remains amenable to updating (Lovibond and Westbrook 2025). If so, the destabilisation of such memories may be especially dependent on the degree of prior learning and the extent to which reminder conditions generate prediction error. Under conditions of asymptotic learning and minimal mismatch, as may have been the case in Cogan et al. (2019), goal-tracking memories may therefore be less likely to destabilise and undergo reconsolidation, even if sign-tracking memories do. This might explain the concurrent impairment of sign-tracking by propranolol in Cogan et al (2019), noting also that this impairment itself contrasts with the previous consistent lack of effect observed with both sucrose (Lee and Everitt 2008a) and ethanol (Milton et al. 2011) rewards.

Another likely reason for the difference between our study and that of Cogan et al ( 2019) is our use of a non-conditioned CS- (note that this also applies to the aforementioned discrepant sign-tracking studies). Our primary outcome is discriminated approach to the CS+ (greater than the CS-)(c.f. Everitt et al. 2001). The use of an explicitly unpaired CS- provides important interpretative value in confirming that behaviour during the CS+ is genuinely a result of the previous CS-US conditioning (Mackintosh 1974). Discriminative approach here was impaired in both reminder conditions; moreover, in the non-reinforced reminder the impaired discrimination manifested as an increase in approach during CS- presentation, with no effect on approach during CS+ presentation. While the behaviour of propranolol-treated rats in the reinforced reminder condition was different (impaired approach during the CS+, with intact approach during the CS-), the observation that impaired discriminative approach can occur in the absence of any meaningful change in approach behaviour during CS+ presentation casts some doubt on whether the findings of Cogan et al (2019) represent a genuine lack of impairment by propranolol.

The second contrasting study was a sign-tracking procedure with an embedded measure of goal-tracking, using discriminated responding to ethanol-associated visual CS+ (vs non-conditioned CS-)(Milton et al. 2011). In this study, MK-801 impaired both sign-tracking and goal-tracking, whereas propranolol did not impair either measure. That the pattern of MK-801 vs propranolol effects on sign-tracking are consistent across sucrose (Lee and Everitt 2008a) and ethanol (Milton et al. 2011) rewards might suggest that the lack of propranolol effect on goal-tracking in Milton et al (2011) is more reliable that the impairment observed in the present study. However, in the Milton et al (2011) study there was seemingly unreliable discriminated approach in the non-reactivation control conditions (at least in the Saline and propranolol groups), and the measure of approach was not normalised against the immediately preceding pre-CS period as in the present study (to account for within-session fluctuations in baseline nosepoking behaviour; c.f. suppression ratios used in forms of aversive conditioning (Mackintosh 1974)).

Although goal-tracking behaviour is often considered less directly translational than sign-tracking, it captures an important and complementary aspect of cue-controlled behaviour: the use of predictive cues to guide actions via expected outcomes. In humans, cue-induced reward seeking frequently depends not only on the motivational properties of cues themselves, but also on learned expectations about forthcoming outcomes, such as the anticipated effects of drugs or the value of food rewards (Milton and Everitt 2012; Lovibond and Westbrook 2025). As such, goal-tracking provides a model of expectancy-based control over behaviour that is highly relevant to maladaptive reward seeking. Consistent with this interpretation, goal-tracking behaviour is sensitive to outcome value and devaluation, indicating a dependence on underlying stimulus–outcome representations that remain amenable to updating (Savage and Ramos 2009; Morrison et al. 2015). Importantly, this capacity for updating makes goal-tracking paradigms particularly well suited for studying reconsolidation-based interventions aimed at modifying maladaptive reward memories, as such interventions are proposed to act on destabilised and labile memory representations following reactivation (Exton-McGuinness et al. 2015; Milton 2023). The present findings therefore suggest that β-adrenergic blockade may be capable of disrupting such expectancy-based processes under appropriate reactivation conditions, highlighting both the translational potential of this approach and the importance of identifying the behavioural and mnemonic boundary conditions that determine the success of reconsolidation-targeted interventions.

In summary, we demonstrate that in a discriminative goal-tracking paradigm, pre-reminder administration of propranolol produces persistent impairments in conditioned responding comparable to those observed with MK-801. These findings challenge previous null results and indicate that β-adrenergic blockade can disrupt reconsolidation of appetitive memories under conditions that promote memory destabilisation. Critically, the effectiveness of this intervention appears to depend on the behavioural and mnemonic parameters of the task, including the strength of prior learning and the nature of the reminder experience. These results therefore support a view of reconsolidation as a boundary-condition-governed process and highlight the importance of identifying the specific conditions under which maladaptive reward memories can be rendered labile and updated. More broadly, they suggest that the translational potential of reconsolidation-based interventions will depend not only on the pharmacological agent employed, but also on the behavioural context in which memory reactivation occurs.

## Data access statement

All data and analyses created during this research are openly available on OSF at https://osf.io/ya4hk/files/osfstorage.

